# Differential replay for reward and punishment paths predicts approach and avoidance

**DOI:** 10.1101/2021.11.18.468950

**Authors:** Jessica McFadyen, Yunzhe Liu, Raymond J Dolan

## Abstract

Planning is thought to involve neural replay, where states relevant to a task goal are rapidly reactivated in sequence. It remains unclear if, during planning, replay of a path relates to an actual prospective choice. Here, using magnetoencephalography (MEG), we studied participants while they planned to either approach or avoid an uncertain environment that contained paths leading to reward and to punishment. We show significant planning-related forward sequential replay with state-to-state transitions in the range of 20 to 90 ms. Replay of rewarding paths was boosted prior to a decision to avoid, and attenuated prior to a decision to approach. Crucially, a trial-by-trial bias in replaying punishing paths predicted an irrational choice to approach when a prospective environment was more risky, an effect that was particularly marked in more anxious participants. The findings reveal a coupling between the content of forwards replay and rational choice behaviour, such that replay prioritises an online representation of potential reward.

## Introduction

When formulating a plan, we are often uncertain as to whether a choice will lead to a good or bad outcome. For example, when we deliberate whether to go to a party or stay home, we might simulate potential series of events that are positive (e.g., arriving and seeing friends, meeting new people, coming home feeling happy) or negative (e.g., arriving and not knowing anybody, saying something embarrassing in front of new people, leaving early and regretting the whole experience). Situations such as these engender an approach-avoidance conflict, wherein decision-making is rendered difficult by uncertainty over whether an action will lead to a positive or negative outcome.

Neural replay, originally characterised in the context of a rapid sequential reactivation of hippocampal place cells that map to specific locations of recently experienced paths ^1–7^, is now linked to a number of functions in both humans and animals, including memory consolidation of spatial ^4,8–11^ and temporal order relationships ^7,12,13^, inference ^14–16^, and credit assignment ^17^. There is also evidence indicating that neural replay may be functionally related to a simulation of potential outcomes during active planning ^18–22^.

A role for replay in planning is supported by observations that when rodents pause during spatial navigation, the order of replayed place cell firing matches paths leading to the learned location of a reward ^23,24^, and is enhanced for paths leading to greater rewards ^25^. Furthermore, the more a rewarding path is replayed, the more likely it is that the animal will correctly pursue that path ^21,23,24,26^. Disruption of replay events at decision points, by application of electric pulses to the hippocampus, leads to the expression of more vicarious trial and error behaviour ^25,27^, and a greater likelihood that an animal will take the wrong path ^10,28^. Remarkably, replay events also provide a mapping of potential trajectories to rewards that have never been experienced, in both online ^29^ and offline ^30,31^ replay.

In contrast to reward, the question of how aversive events modulate replay is relatively under-investigated. Animal studies show that removal of a reward leads to a marked reduction in replay ^32^. Paths leading to danger are also more strongly replayed, and this is anticorrelated with an animal’s chosen trajectory, such that an animal tends to avoid the dangerous path ^33^. Such findings have led to a proposal that hippocampal replay prioritises paths that are most immediately relevant for on-going behaviour ^34^.

A feature of previous studies is that they tend to use environments that contain either reward or punishment. How replay is impacted in a prospective environment where paths might lead to either reward or punishment is unknown, where such environments give rise to an approach-avoidance conflict ^35,36^. An inability to make optimal decisions under approach-avoidance conflict is a characteristic of clinical anxiety disorders, where the potential for experiencing a negative event leads to avoidance regardless of the likelihood of potential reward ^37–39^. On the other hand, a tendency to approach, even when dangerous to do so, is considered a risk factor for developing substance abuse disorders ^40^. During approach-avoidance conflict, interactions between anterior and ventral hippocampus are thought to arbitrate decisions to approach or avoid in both humans ^36,41^ and animals ^42,43^ via a monitoring of the magnitude and likelihood of threat. Replay is a potential candidate mechanism for this process, where a relative increase in replay strength of one trajectory over another could, in principle, relate to a bias towards approach or avoidance.

In this experiment, we employed magnetoencephalography (MEG) to investigate whether there is an asymmetry between the replay of rewarding and aversive sequences during planning. We designed a gambling-style task in which participants made decisions to either approach or avoid an uncertain environment containing gains and losses, with the goal of earning as many points as possible. By decoding rapid sequential replay related to sequences of transitioned states, we reveal a striking asymmetry in replay that reflects prospective evaluations during planning and predicts trial-by-trial decision-making.

## Results

### Expected value guides decision-making

Participants learnt two sequences (hereafter referred to as “paths”) each containing three images (hereafter referred to as “states”) with an associated integer value (**Supp. Fig. 1C**). In a gambling-type scenario, participants could choose to either “approach” an environment containing the two paths (probabilistically transitioning to one of the two paths) or “avoid” this environment entirely (meaning they would receive a guaranteed sum of 1 point), where the overall task goal was to earn as many points as possible.

**Figure 1.**
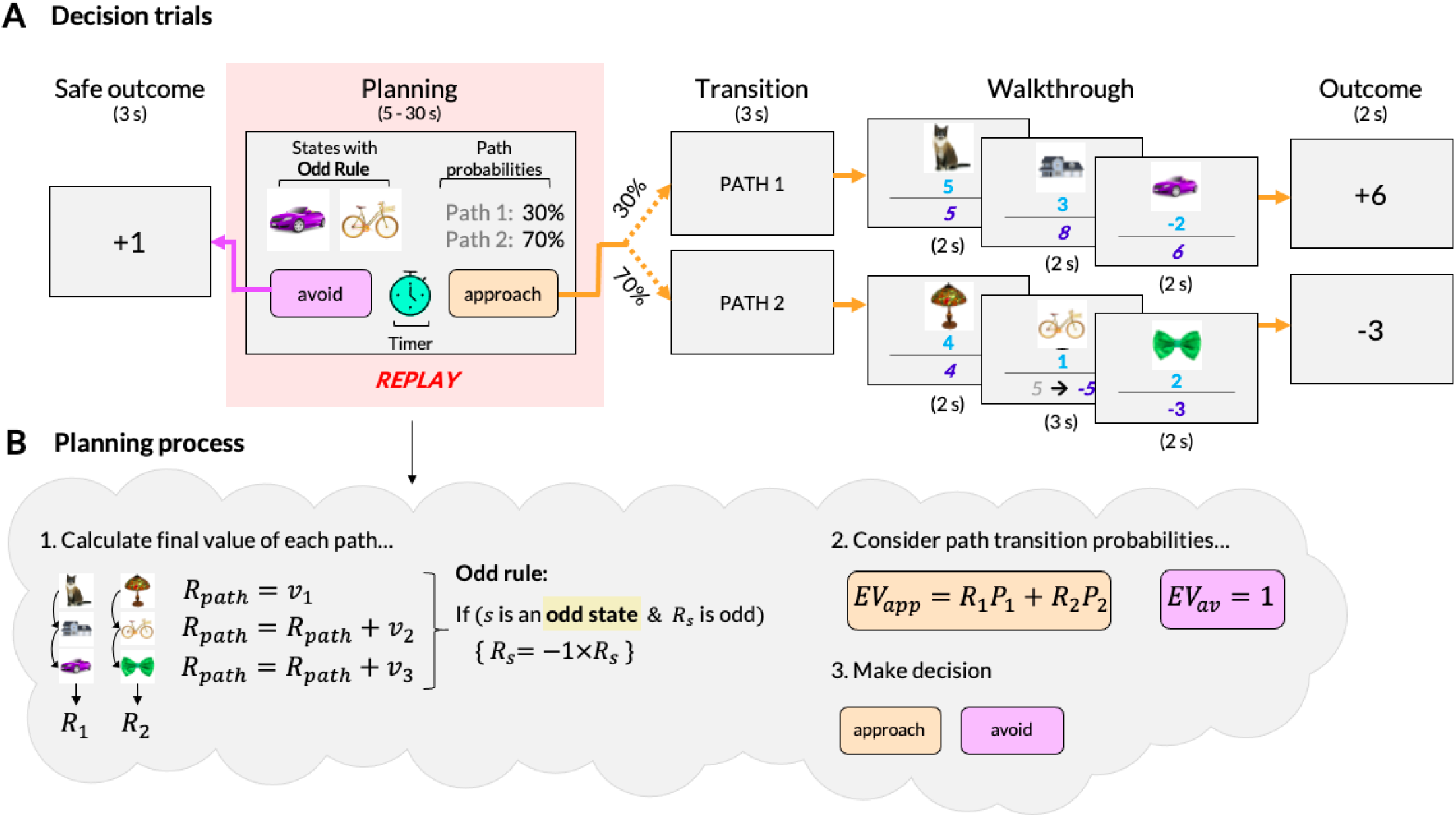
Decision trials. (**A**) Participants began each trial in a planning phase. Here, they used the presented information (the odd rule states and the path transition probabilities) to mentally calculate the total outcome for each path and evaluate the utility of an approach vs avoid decision. This calculation involves summing the value (*v*) of each state (*s*) across each path, taking into account the ‘odd rule’, and multiplying the final sum (***R***_***path***_) by the path transition probabilities (***P***_***path***_), as described in **B.** The odd rule encouraged conscious sequential planning along task states, and discouraged employment of a deterministic strategy. The order of images and their respective values were learned in an initial training phase (see **Supp. Fig. 1**). MEG data from this planning period was the focus of replay analysis. If participants chose to approach, a screen appeared displaying which of two potential paths they had transitioned to (“Transition” screen), and participants then observed an animation of the sequence (“Walkthrough” screens). During this walkthrough, the number of points gained or lost in each state (light blue numbers), and the cumulative sum (dark blue numbers), was shown below the state image. Note that images were only shown in forced-choice trials, while text labels were shown in all other trials. The final sum of points for the sequence was then shown (“Outcome” screen). If participants chose to avoid, an increase of one point was shown (“Safe outcome” screen). (**B**) The odd rule was introduced to encourage prospective planning and entailed that, if the cumulative sum of points collected up to (and including) an odd rule state was an uneven number, then the sign of the cumulative sum up to and including that state would be flipped (e.g., −3 would become 3, or vice versa), and this new amount carried over to be added to the value of any remaining states. In each trial, the odd rule was applied to one state from each of the two prospective paths. A rational planner first calculates the cumulative sum of points along each path (taking the odd rule into account), multiplies these by the respective path transition probabilities (which varied trial to trial), and on this basis then decides based upon the expected value whether to approach (***EV***_***app***_) or avoid (***EV***_***av***_).

To make an informed choice, participants needed to mentally simulate an accumulation of points along each path. The total value of each path was dependent on a visual cue presented at the beginning of each trial (**Fig. 1**; see description of “odd rule” in **Methods**). In a majority of trials, one of the two paths resulted in a points gain and the other in a loss. The probability of transitioning to either path (if participants chose to approach) spanned five probabilities (10-90%, 30-70%, 50-50%, 70-30%, and 90-10%) displayed on screen during the 30-second planning period.

If participants chose to approach, a screen then displayed which of the two available paths had been selected, where this was determined by the path transition probabilities displayed during the planning phase (**Fig. 1A**). Participants then deterministically transitioned to each state along the selected path, with the state value and cumulative sum of points along the trajectory displayed underneath. Note that the first four trials of each block were forced-choice to approach. This provided participants with a reminder of the images representing each state (images were replaced by text labels in all other free-choice trials to control visual exposure) and the associated integer values (the value of one state from each of the two paths was updated at the beginning of each block). If participants chose to avoid, a screen was displayed stating that participants had earned one point.

In the task, rational decision-making entails calculating the expected value of approaching (i.e., the sum of points for paths 1 and 2, weighted by their probabilities) and choosing to approach only if the expected value is greater than the certain value available by choosing to avoid (i.e., ≥ 1). We calculated the accuracy of participants’ choices by comparing them to rational choice behaviour in the task. Two of 26 participants performed with only 47.55% and 51.37% accuracy and thus were excluded from all subsequent analyses, except for evaluation of replay of an overall state map.

Participants performed significantly above chance, with 76.07% accuracy on average (SD = 7.35%, range = 60.27% to 89.73%; t(23) = 17.373, p < 0.001; **Fig. 2A**), correctly approaching when the expected value of approaching was 2.386 on average (SD = 0.57) and avoiding when the expected value was −1.552 on average (SD = 0.556; t(23) = 21.152, p < 0.001; **Fig. 2B**). Overall, participants tended to approach more often (57.15% of trials) than avoid (42.85%; t(23) = 4.176, p < 0.001), consistent with reward-seeking or information-seeking behaviour. Experimental protocols were designed so that the expected value of approaching was > 1 on 50% of trials (**Supp. Fig**.**1E**). As such, accuracy was significantly lower on trials where participants chose to approach (74.59%, SD = 7.53%) than to avoid (79.27%, SD = 8.07%; t(23) = −3.190, p = 0.004; **Fig. 2A**), as expected for reward-seeking or information-seeking behaviour. Participants were also significantly faster to approach (M = 8.446 seconds, SD = 2.03) than to avoid (M = 8.975 seconds, SD = 2.424; t(23) = −2.319, p = 0.030; **Fig. 2C**).

**Figure 2.**
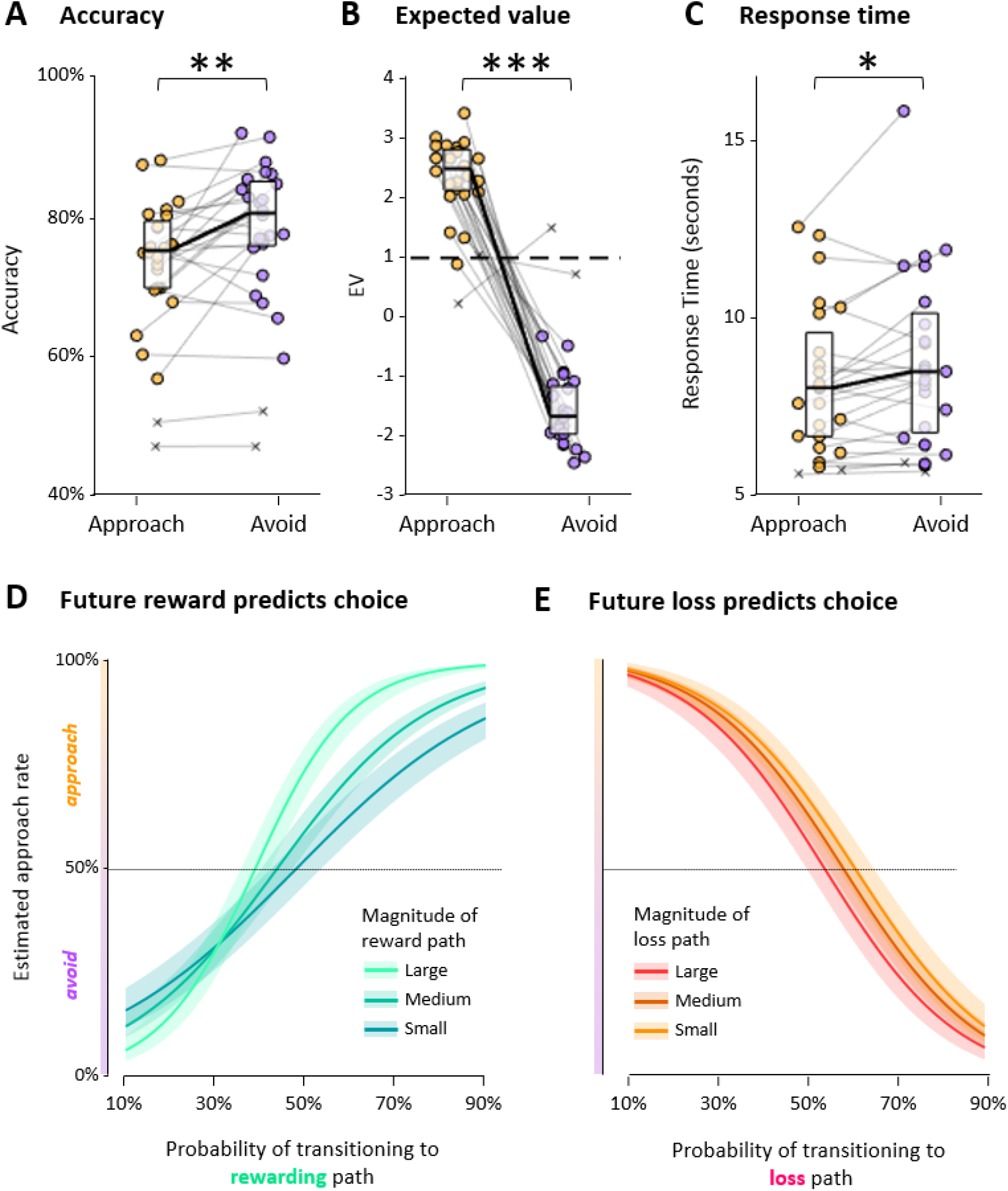
Behavioural results. **(A)** Accuracy is the proportion of trials in which participant responses matched an optimal response, based upon expected value. Overall, participants (individual markers) made significantly more accurate avoid (purple) decisions than approach (orange) decisions. **(B)** The expected value of approaching when participants chose to approach was significantly higher than when participants chose to avoid. **(C)** Participants were significantly faster to approach than to avoid. Boxplots indicate median and 25th and 75th percentiles of average participant response times. **(D)** Approach rate estimated by a behavioural multilevel model. This shows that participants were more likely to approach if the probability of transitioning to a rewarding path was higher, especially when prospective reward values were greater. **(E)** Similarly, participants were more likely to approach if potential loss was lower, irrespective of path transition probability. * p < .05, ** p < .01, *** p < 0.001.

In the experimental design, there was consistency as to which of the two paths culminated in a reward or loss. Hence, for the first half of the experiment, path 1 resulted in reward and path 2 resulted in loss, and vice versa for the second half of the experiment (two protocols were used and counterbalanced across participants; see **Methods**). To encourage active engagement in sequential planning, rather than merely learning this tendency, we included catch trials (5%) where either both paths lead to a reward or both lead to a loss. Although accuracy was numerically poorer on catch (67.41%) compared to regular trials (74.54%), this difference was not statistically significant (t(23) = 1.910, p = 0.069). Furthermore, accuracy did not significantly differ between the last block before a contingency swap and the first block of the contingency swap (block 5 = 71.43%, block 6 = 73.80%; t(23) = −0.700, p = 0.491). Altogether, these results indicate participants engaged in online evaluation of each path during planning, and incorporated trial-specific transition probabilities into their decision-making.

We next constructed a multilevel logistic regression model to more precisely examine how path values and transition probabilities influenced trial-by-trial decision-making. In our model, trial-by-trial choice was predicted by a three-way interaction between the value of the path with the highest prospective value (i.e., the rewarding path), the value of the path with the lowest prospective value (i.e., the loss path), and the probability of transitioning to the rewarding path (note that the probability of transitioning to one path was always relative to the other). We also included response time as a fixed effect, as well as certainty of path transition probabilities on each trial (uncertain: 50-50%, moderately certain: 30-70% or 70-30%, very certain: 10-90% or 90-10%).

Approach choices were significantly predicted both by the probability of transitioning to a more rewarding path (β = 6.460, p < 0.001) and by larger prospective rewards (β = 0.113, p < 0.001; **Fig. 1F**). Thus, participants approached environments containing larger rewards more so when the probability of transitioning to reward was higher (β = 4.991, p < 0.001). Although the magnitude of potential loss also predicted decision-making (participants were more likely to choose to avoid when potential losses were larger: β = 0.053, p = 0.011), there was no interaction with transition probability (β = −0.028, p = 0.744; **Fig. 1G**). These findings support the idea that decision-making was guided by total value of reward and loss paths, as well as the probability of transitioning to a rewarding path. We also observed a significant effect of certainty (β = 0.317, p < 0.001), such that participants were more likely to approach when transition probabilities were more certain overall (i.e., 90-10% or 10-90%, as opposed to 50-50%). Finally, given that participants were more likely to approach when rewarding paths were more probable, we tested whether participants experienced rewarding paths more frequently than aversive paths. On average, participants transitioned to a rewarding path 107 times (SD = 11) and to an aversive path 23 times (SD = 8), a difference that was significant (t(23) = 25.577, p < 0.001). There was no significant difference in the likelihood of experiencing path 1 (M = 46, SD = 8) or path 2 (M = 46, SD = 7; t(23) = −0.046, p = 0.964) due to our counterbalanced design.

### Forward replay during planning

Our primary research questions were: 1) whether there is replay of state-to-state transitions during planning, 2) whether this is influenced by prospective value calculations of each path’s value, and 3) whether, in turn, this relates to a subsequent choice to approach or avoid. In an initial functional localiser task, we trained classifiers on visually-evoked response fields, measured using MEG, to each of the six unique state images (see **Fig. 3A-C**). Importantly, these state representations were captured prior to participants learning the order of states in each sequence. We then applied each state classifier to MEG data acquired during the planning period of each decision trial, producing time series of decoded state reactivation (**Fig. 3D**). We used general linear modelling to assess evidence for temporally-ordered reactivation of each state pair (A-B and B-C in path 1, and D-E, and E-F in path 2) across different time intervals (10 to 600 ms, in steps of 10 ms), in both a forwards and backwards direction. We refer to this as “sequenceness”, our index of replay.

**Figure 3.**
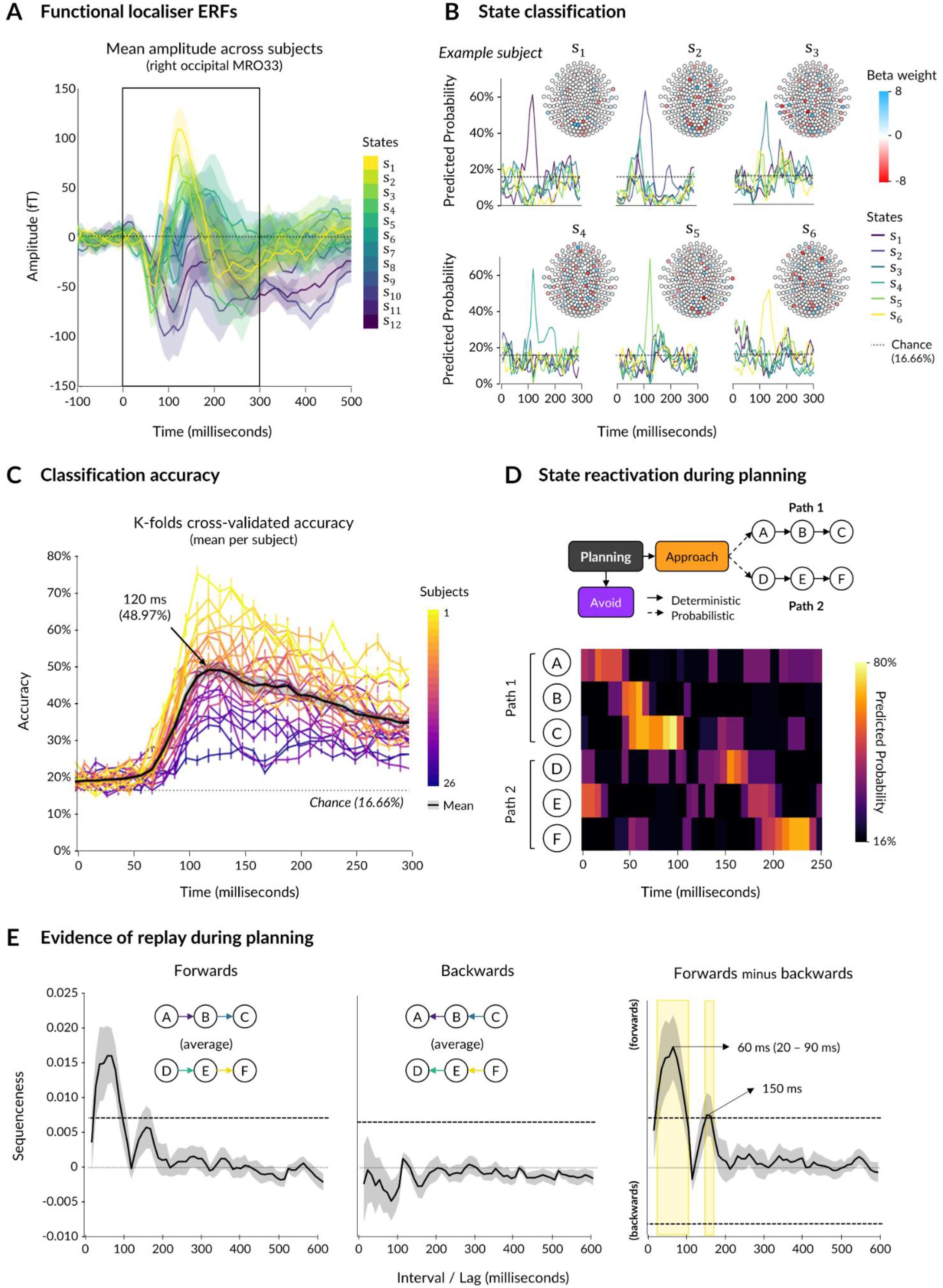
State classification and replay analysis. **(A)** Before learning the order of images along each path, participants viewed each image in an initial functional localiser task. The visually-evoked event-related fields (measured using MEG) are displayed for each of the 12 images, or “states” (6 were randomly assigned to each participant), averaged across participants. **(B)** Using the functional localiser MEG data, classifiers were created for each state, per participant (example participant is shown). A classifier was a set of beta weights per sensor. **(C)** Using K-folds cross-validation, we assessed average accuracy across state classifiers per participant. Classifiers trained at a 120 ms time point produced the highest average accuracy overall. **(D)** Classifiers trained on either 110, 120, or 130 ms were applied to MEG data collected throughout the planning period of decision trials, producing matrices of predicted state reactivation per trial (example shown). **(E)** Using a two-level GLM approach, we estimated the intervals (or “lags”) between maximal reactivation of each state during planning, in a forwards (left) and backwards (middle) direction. Plots display the sequenceness estimates averaged across all four transitions. The significance threshold is indicated by an horizontal dashed line. Significant forwards-minus-backwards replay occurred at state-to-state intervals of 20 to 90 ms, peaking at 60 ms with a second, albeit less pronounced, peak at 150ms.

As a first step, we asked whether there was evidence for replay of the entire state space (i.e., average sequenceness across all four transitions), discarding the first four trials in each block as these were forced-choice. We observed maximal forward state-to-state reactivation at 60 ms intervals (or “lags”), and maximal backward state-to-state reactivation at 110 ms (**Fig. 3E**). We then computed a forward-minus-backward sequenceness measure to remove common noise and increase sensitivity. A significance threshold generated by random permutations (see **Methods**) provided evidence for significant forward replay at both 20 to 90 ms and 150 ms state-to-state intervals. These results indicate that, during planning, the state space was replayed with a rapid temporal compression akin to that reported in previous studies ^12,14,15,17,44^.

### Replay is modulated by prospective reward and loss

Having found evidence for forwards replay during planning, we next asked whether we could differentiate replay for paths that culminated in either a reward or a loss. For each trial, we averaged sequenceness across the two transitions in each path (**Fig. 4A**). We then entered these trial-by-trial estimates of path replay at the significant state-to-state intervals identified by our previous analysis (20 to 90 ms) into a linear mixed-effects model, accounting for effects of subject, replay interval, and trial duration. We then modelled whether path type (i.e., reward or loss) predicted replay strength during planning, and whether this interacted with post-planning choice.

**Figure 4.**
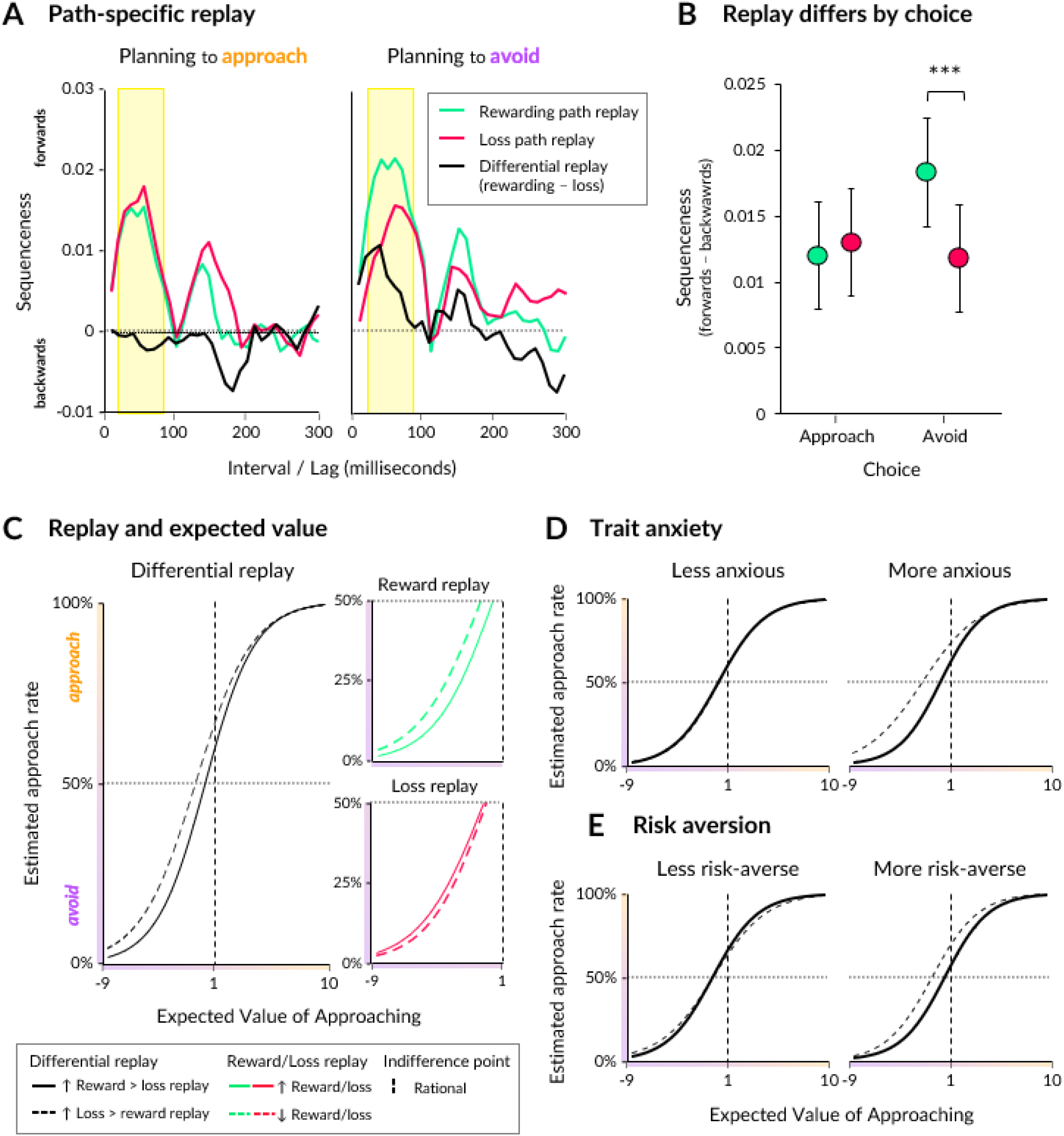
Replay of prospective reward and loss paths. **(A)** Replay strength for paths leading to either reward (green) or loss (red), split according to whether participants chose to approach (left) or avoid (right). Data is averaged across trials and participants. Significant replay intervals are highlighted by the yellow box. The difference between reward and loss replay is also shown (black). **(B)** Estimated marginal means produced by a mixed-effects model predicting replay strength by the total value of a path (reward in green, loss in red) and the choice subsequently made by participants (approach or avoid). Error bars indicate 95% confidence interval. * p < .05, ** p < .01, *** p < 0.001 **(C)** The approach rate (y axis) predicted by a mixed-effects model containing expected value (x axis) and differential replay. When differential replay was more negative (dashed line; indicates relatively stronger replay of loss than reward paths), participants were more likely to approach environments with lower prospects (i.e., negative expected value). A similar model using separate predictors for reward and loss path replay showed participants were more likely to approach negative expected value trials when replay of rewarding paths was diminished (green dashed) and when replay of loss paths was enhanced (red solid). The indifference point (i.e., the point at which approach rate should be 50%) is displayed for rational behaviour (vertical dashed line). **(D)** Same as **C**, except that participants’ trait anxiety and risk-aversion scores were included as interactors in the model. The interaction between differential replay and expected value on choice was driven predominantly by more anxious participants (low/high split is for visualisation purposes only). **(E)** Same as **D**, except data has been split into low and high risk-aversion. More risk-averse participants were more likely to approach when differential replay was more negative, regardless of expected value.

Overall, rewarding paths were replayed more strongly than aversive paths (β = 0.007, p < 0.001; **Fig. 4B**). A significant interaction with choice (β = −0.008, p < 0.001) revealed that this stronger replay for rewarding paths occurred primarily when participants planned to avoid (EMM_Δ_ = −0.007, p < 0.001), while replay strength did not differ between reward and loss paths when participants planned to approach (EMM_Δ_ = 0.001, p = .762). Thus, path replay strength was conditional on subsequent choice, such that replay of a rewarding path was modulated by a current instrumental goal (i.e., to obtain reward or avoid punishment).

We next performed two analyses designed to test whether replay was modulated by factors other than choice; namely, recent path experience, and the probability of transitioning to either path irrespective of path value. First, we asked whether recent path experience influences replay of prospective paths. We operationalised path experience as the number of trials within a block since participants last experienced a particular path. Applying a log-transformation to address a positive skew, we added this as a predictor interacting with path type (reward or loss) and transition probability. Intriguingly, we found an interaction between path type and experience (β = 0.010, p < 0.001). This showed that reward paths were more strongly replayed when they had not been experienced for longer intervals, whereas loss paths were more weakly replayed (**Supp. Fig. 2A**). The probability of transitioning to each path modulated this effect (β = 0.014, p = 0.010), such that less recently-experienced rewarding paths were even more strongly replayed when the upcoming transition probability was higher. Thus, recent experience had opposing effects on the replay of reward and loss paths.

Second, we asked whether path replay, irrespective of reward or loss, was modulated by transition probability. We modelled replay of each path per trial as being predicted by each path’s transition probability, as well as the subsequent choice made on each trial. We found that replay was significantly stronger overall when participants planned to avoid (β = −0.003, p < 0.001), but found no significant effect of the probability of transitioning to the approach paths (β < 0.001, p = 0.830) nor any interaction with choice (β = −0.004, p = 0.190; **Supp. Fig. 2B**). This indicates that participants’ beliefs about which path is more likely to be experienced did not impact the strength of replay.

### Replay predicts approach and avoidance

The stronger replay for rewarding paths when subjects planned to avoid points to a relationship between the content of replay and decision-making. To investigate this further, we computed a measure of “differential” replay that captures a difference in the expression of sequenceness between each prospective path on a trial-by-trial basis. Specifically, we subtracted loss path replay from reward path replay, such that more positive differential replay indicates a bias towards replaying paths leading to reward, and vice versa for more negative differential replay.

Using this differential replay measure, we next modelled how replay content changed conditional on the choice being planned (i.e., to approach or avoid), as well as the environment prospects (i.e., the cumulative gain or loss for each path, and the probability of transitioning to each path). To simplify the model, we used the expected value of approaching on each trial as a summary metric of an environment’s prospects (equivalent to the total sum of points for each path weighted by their respective transition probabilities). For predicting trial-by-trial decision-making, we constructed the model to allow expected value to interact with differential replay at all significant replay intervals (20 to 90 ms), as well as certainty of path transition probabilities and response times.

At a behavioural level, there was a sigmoidal relationship between expected value and choice (β = 0.432, p < 0.001; **Fig. 4C**), such that participants were more likely to approach when the associated expected value was ≥ −1.2. This is below a rational indifference point of 1, indicating participants were more likely to approach environments with poorer prospects. Additionally, participants were more likely to approach when path transition probabilities were more certain (β = 0.270, p < 0.001).

At a neural level, trial-by-trial differential neural replay predicted choice (β = −0.713, p < 0.001), such that participants were more likely to approach when differential replay during planning was less positive, reflecting a bias towards replaying paths leading to potential loss and/or a bias away from replaying paths leading to potential reward. Importantly, this effect of differential replay on decision-making interacted with expected value (β = 0.133, p = 0.008), such that a bias away from replaying paths leading to reward was even more pronounced when participants planned to approach environments with a more negative expected value.

Our use of a difference measure precluded knowing whether the above effect was driven by diminished replay of reward paths or enhanced replay of loss paths. To unpack this further, we duplicated our model but now replaced differential replay with two separate predictors for reward path replay and loss path replay, each separately interacting with expected value. This revealed that path replay for reward and loss had opposing interactions with expected value. Planning to approach a more hazardous environment (i.e., negative expected value) was predicted by enhanced replay of paths leading to loss (β = 0.120, p = 0.090) and diminished replay of paths leading to reward (β = −0.146, p = 0.031; **Fig. 4C**). Moreover, as suggested by our initial analyses on replay and path value, when participants planned to approach, replay of reward paths was significantly reduced (β = −1.232, p < 0.001). Replay of loss paths did not predict decision-making overall (β = 0.189, p = 0.313). Thus, overall, the content of replay during planning predicted subsequent decisions such that, when paths leading to reward were selectively replayed, participants exhibited more rational decision-making by choosing to avoid when expected value was lower. In contrast, when replay of reward paths was reduced (leaving relatively stronger replay of loss paths), participants were more likely to approach riskier environments.

### Trait anxiety and risk aversion

Next, we tested an hypothesis that the relationship between differential replay and approaching risky environments is amplified in participants with higher trait anxiety or a higher propensity towards risk-taking. To assess this, we performed an independent components analysis on self-report questionnaires, yielding one component representing anxiety and another representing risk-aversion (see **Methods**). We then constructed a model in which each of these personality traits interacted with both differential replay during planning and expected value to predict future decision-making. Again, the degree of certainty about the path transition probabilities was included in the model, as well as trial duration.

Within this model, anxiety and risk-aversion alone did not predict decision-making (β = −0.028, p = 0.644 and β = 0.073, p = 0.284, respectively), although there was a significant increase in approach rate at higher expected values (indicating more conservative decision-making) in participants with higher risk aversion (β = 0.008, p = 0.048). Instead, both anxiety and risk aversion significantly modulated the relationship between differential replay and decision-making. More anxious participants (β = −0.314, p = 0.003; **Fig. 4D**) and more risk-averse participants (β = −0.377, p < 0.001; **Fig. 4E**) were more likely to approach when replay was biased away from rewarding paths (β = −0.314, p = 0.003). For more risk-averse participants, this occurred regardless of expected value (β = −0.018, p = 0.540), whereas for more anxious participants, this occurred predominantly when expected value was lower (β = 0.096, p = 0.014). Thus, the previous finding that replay is biased away from rewarding paths when planning to approach more precarious environments was driven by more anxious participants.

We repeated this model using separate interacting predictors for reward and loss path replay to discern which was driving the effects described above. The model revealed that replay for paths leading to loss (β = 0.450, p = 0.003), but not reward(β = −0.199, p = 0.189), was boosted for more anxious participants when approaching more aversive environments. Similarly, replay for paths leading to loss (β = 0.483, p < 0.001), but not reward (β = −0.225, p = 0.084), was boosted for more risk-averse participants when planning to approach any environment. Additionally, more risk-averse participants had diminished replay of rewarding paths when planning to approach more aversive environments, while more risk-seeking participants had diminished replay of rewarding paths when planning to approach more lucrative environments (β = −0.132, p = 0.004). In contrast, more anxious participants had stronger replay of loss paths when planning to approach more aversive environments (β = −0.201, p < 0.001). This suggests that the more negative differential replay in participants with higher trait anxiety during planning was driven by an increase in loss path replay rather than a decrease in reward path replay, while the opposite was true for approach planning in more risk-averse participants.

### A prefrontal-temporal network for replay

In our final analysis, we estimated the spatial sources underlying onset of replay events for rewarding and punishing paths. We defined a replay “event” as maximal reactivation of one state followed by maximal reactivation of the following state within a 20 to 90 ms lag, with additional stringent criteria (see **Methods**). First, we determined the underlying sources for any replay event for any transition, regardless of reward or loss, focusing on the left and right hippocampus as regions of interest. Applying a small volume correction, we found a significant increase in broadband (1 to 150 Hz) power in both left (MNI: 40, −16, −17, k = 49, p = 0.009 FWE-corrected) and right (MNI: −35 −21 −12, k = 30, p = 0.009 FWE-corrected) hippocampus at onset of replay events, akin to previous studies using MEG to measure replay in humans ^14,15,44^.

When we expanded our analysis to the whole brain, we observed significant power increases in the frontal regions (including prefrontal and orbital-frontal cortices) and temporal cortex (including superior temporal gyrus and temporal pole; **Fig. 5**). We also observed significant clusters in the mid-occipital cortex, thalamus, insula, cingulate gyrus, and pre- and post-central gyri (whole brain, p < 0.05 FWE-corrected). These results align with previous studies that also report replay in the prefrontal cortex during planning ^45–47^, as well as studies using MEG that also report clusters in the occipital and parietal cortices ^15,44^. See **Supplementary materials** for further source localisation of replay events for reward and loss paths.

**Figure 5.**
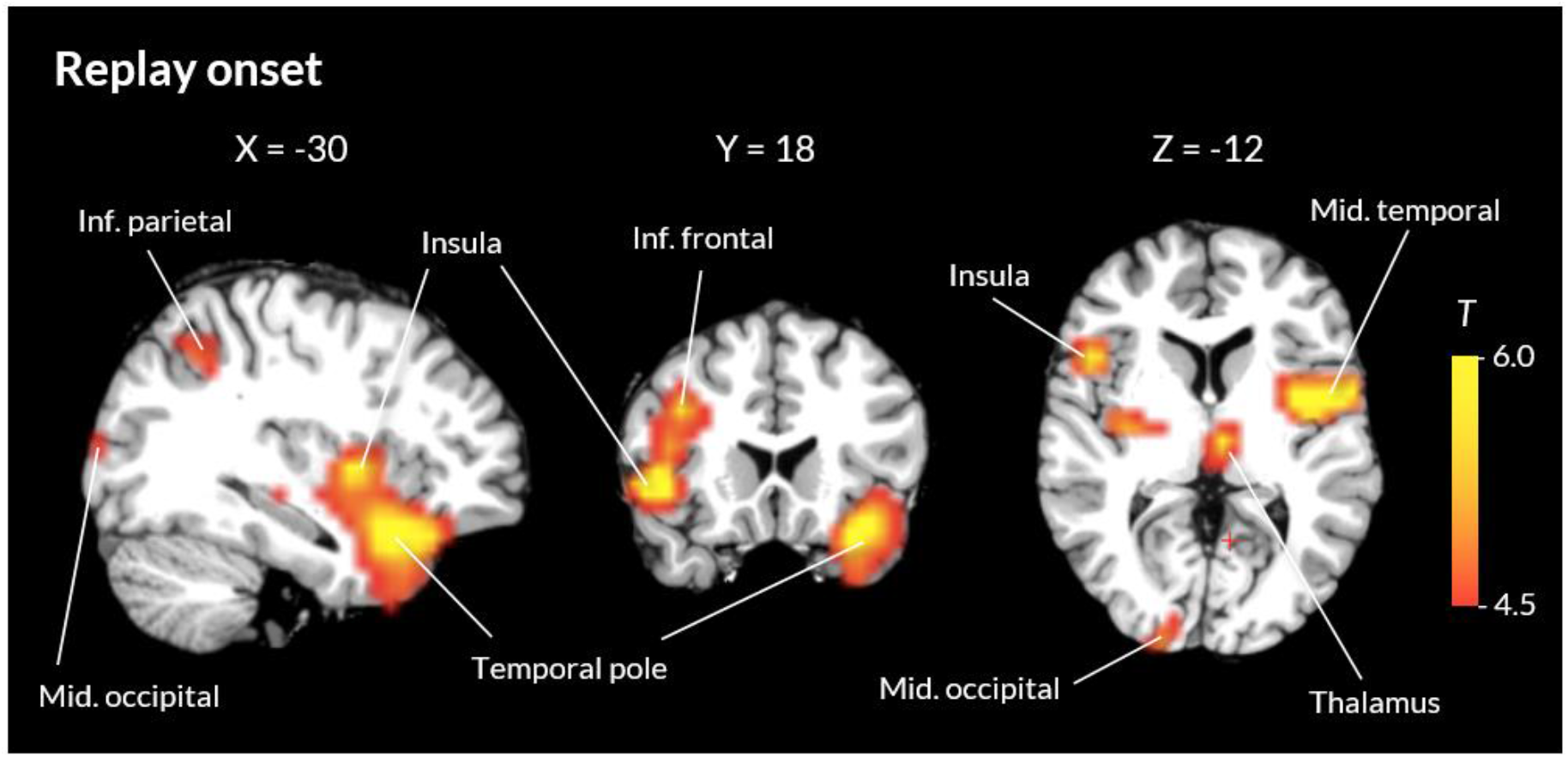
Beamforming analysis on replay onsets. Sources underlying the onset of replay events for any state-to-state transition included the frontal cortex, temporal cortex, insula, and thalamus. Clusters are thresholded at p < 0.05, whole brain FWE-corrected.

## Discussion

In rodent studies, replay content during planning reflects paths that should be pursued ^23,24^ as well as those that should be avoided ^33^. Here, in human subjects, we show that the content of forward replay during planning flexibly predicts decision-making in the context of an approach-avoidance conflict. Participants were more likely to avoid when replay was relatively stronger for paths leading to reward, and more likely to approach when replay was relatively stronger for paths leading to loss, an effect most pronounced when environments were more hazardous (i.e., negative expected value of approaching). This effect was amplified in participants with higher trait anxiety. Our findings support a role for replay in planning under uncertainty, where the relative strength between reward and loss path replay predicted whether decision-making was rational or irrational.

Based on rodent studies, we expected replay content to reflect the different goals of approach (to obtain reward) and avoidance (to avoid punishment), such that replay increases for rewarding paths being pursued ^21,23,24,26,48^ and for punishing paths being avoided ^33^. Instead, we observed the opposite. Preceding a decision to avoid, replay was increased for paths leading to a foregone reward. In contrast, replay of paths leading to reward was decreased preceding a decision to approach. In fact, when there was greater risk associated with approaching (i.e., negative expected value), replay increased for paths leading to potential loss. This pattern suggests a relationship between the content of replay and rational decision-making, with stronger replay of rewarding paths predicting rational avoidance, while a shift towards replaying punishing paths predicts irrational approach.

The above findings were most pronounced in participants who self-reported higher anxiety (specifically, an intolerance of uncertainty and a propensity to worry), who replayed potential punishing paths to a greater extent when planning to approach aversive environments. Similarly, for participants who self-reported higher trait risk-aversion, replay was biased away from reward and towards loss when planning to approach, regardless of the expected value of approaching. Thus, trial-by-trial instances of irrational approach-avoidance behaviour were more strongly related to enhanced replay of potential loss paths in more anxious and more risk-averse participants.

Replay has been speculated to play a role in clinical anxiety and depression ^38^. Our study is the first to provide evidence for a relationship between replay and trait anxiety, as well as risk-aversion. Dispositional anxiety is associated with a heightened, and sometimes uncontrollable, simulation of potential past (rumination) or future (worry) aversive events ^37,49^. Given replay may play a functional role in selectively sampling a prospective environment during planning, replay might plausibly be biased towards more aversive outcomes in people who have a greater tendency to worry ^37^. Indeed, people with higher social anxiety engage in “counterfactual” updating, entailing stronger deliberation of outcomes that have not, or will not, be experienced ^50^. A simulation of “what if” scenarios maps closely onto our finding that replay content reflects a worst-case scenario of a plan to approach (i.e. the possibility of being punished) or avoid (i.e., forgoing potential reward).

It is important to note that more anxious and more risk-averse participants did not make more erroneous approach decisions overall. Thus, the effect of anxiety and risk-aversion on decision-making was only discernible from underlying neural activity. Our sample consisted of healthy controls recruited from the general public. Future studies involving participants with clinical anxiety disorders who show irrational risky choice behaviour ^51^ will help elucidate the extent to which replay relates to anxiety-modulated decision-making.

In previous studies, sequences leading to reward and loss are typically learned from experience. As such, reinforcement learning is often used as a framework for understanding the mechanistic role of replay in decision-making ^17,34,52–55^. A novel aspect of our study is that, rather than learning from experience, reward and loss were calculated prospectively on each trial by using a combination of visual cues, mathematical calculations, and a pre-learned temporal structure. Additionally, the inclusion of catch trials discouraged learning from experience. As such, it is difficult to relate our findings to a normative account suggesting replay subserves planning by prioritising access to memories for sequences that have greater utility ^34^. Instead, our results suggest that replay prioritises a simulation of outcomes that would *hypothetically* instantiate a change in policy (i.e., a worst-case scenario of approaching and avoiding), despite subjects having no direct experience of these specific outcomes.

Replay of hypothetical or inferred sequences has previously been explained by a role for replay in the maintenance of cognitive maps ^14,15,18,29–31,56^. Similarly, replay content that is decoupled from (or counterfactual to) subsequent choice behaviour has been explained by a role for replay in preventing cognitive maps from becoming biased by an over-representation of repeatedly chosen paths, thus ensuring flexible decision-making in the future ^22,57,58^. A role for replay in memory maintenance does not comprehensively explain our observation of a choice-related modulation of replay in the absence of learning. Such an account would predict increased replay for both paths when planning to avoid (as neither will be experienced), as well as increased replay for the less probable path on each trial when planning to approach. While we observed an overall increase in replay preceding avoidance, it was rewarding paths alone that were more strongly replayed. We also found that rewarding paths were replayed more when they had been less recently experienced, supporting previous research ^58^, but this effect was amplified when the probability of transitioning to the rewarding path was higher, rather than lower. Moreover, punishing paths were replayed to a lesser extent when they had been less recently experienced. Thus, our findings for rewarding path replay partially support a memory maintenance account, while our findings for punishing path replay are more difficult to reconcile ^57,58^.

A possibility is that, in our study, reward was perceived by participants as more salient than loss, in line with participants’ choices being more sensitive to probability and magnitude of reward than that of loss. Playing to accumulate monetary rewards, as opposed to accumulating monetary losses, has been shown to enhance the utility of reward ^59^. This is a potential explanation for why replay reflected a worst-case scenario of choosing to avoid (i.e., forgoing potential reward) across all trials irrespective of expected value, whereas replay reflected a worst-case scenario of choosing to approach (i.e., transitioning to a loss path) only when the expected value of approaching was more negative. Employing a variant of the current design using more arousing positive and negative stimuli (e.g., electric shocks or affective visual stimuli) might provide further insight into the aforementioned possibilities.

Our study presents novel evidence for a relationship between replay content and decision-making that is unrelated to learning and is not fully explained by recent experience. Furthermore, we show evidence for a pivotal role of trait anxiety, such that path prioritisation for replay reflected a worst-case scenario of a decision to approach (increased replay for loss paths) or avoid (increased replay for reward paths). These results indicate current theories of replay might need to accommodate situations in which direct experience of a sequence and its outcome is unavailable. This scenario is particularly pertinent to survival, where an outcome might include the death of the agent ^60^, as well as to our understanding of anxiety disorders, which are characterised by an over-simulation of improbable but catastrophic events ^37^.

## Methods

### Participants

We recruited 32 healthy volunteers via online advertisements to participate in the first session, which served as an opportunity to practice and as a screening point to exclude participants who found the memorisation or arithmetic in the task too difficult (see **Methods, Experimental task**). We excluded 1 participant who scored < 80% accuracy when tested on the image order, and 4 participants who scored < 60% accuracy in the decision trials. Thus, 27 participants completed session 2. One of these participants was excluded due to a technical error with MEG data collection. The final sample consisted of 26 right-handed participants (8 males, 18 females) aged between 18 and 35 years (M = 25, SD = 5).

All participants were fluent or native English speakers with normal vision and no current use of psychiatric medication. The study was approved by the University College London Research Ethics Committee (9929/002). Each participant provided written consent for each session and were paid £50 (£10 for behavioural session and £40 for MEG session), plus up to £15 bonus (up to £5 for the behavioural session and up to £10 for the MEG session) upon completing the study. Bonuses were calculated by converting the accuracy of each block (i.e., the proportion of times participants made the correct choice) into a monetary value between £0 and £1.

### Experimental task

#### Image learning

The experiment was created for web browser using jsPsych v6.1.0 31. The experiment was presented in the format of a computer game where participants played the role of an astronaut exploring rooms within a spaceship. There were six rooms in total, arranged as two sequences (or “paths”): path 1 contained rooms A, B, and C, and path 2 contained rooms D, E, and F. Each room (or “state”) was represented by a unique image randomly selected from a set of 12 for each participant (**Supp. Fig. 1A**). During the image learning phase, participants watched an animation of the transitions along each path, in which the images for each room were presented one at a time for 3 seconds each (**Supp. Fig. 1C**). Participants were then tested on their memory for the order of images in each sequence. Participants were given up to two attempts to reach at least 80% accuracy.

#### Value learning

After successfully completing the image learning phase, participants then learned to associate an integer value (ranging from −5 to 5, excluding 0) with each room. This integer represented the number of points subjects stood to gain or lose in each room. To learn these values, participants were presented with each sequence four times, with the integer value presented underneath each image (4-second presentation; **Supp. Fig. 1D**). Participants were then tested on their memory for each individual room’s value, as well as their ability to calculate the cumulative sum of points in each room. This process was repeated until participants scored at least 80% accuracy (up to two attempts).

#### Decision trials

After completing image and value learning, participants then partook in decision trials. At the beginning of each decision trial, participants were placed conceptually “outside” of the environment containing the two learned sequences and could choose to either approach or avoid it (**Fig. 1A**). Avoidance resulted in a guaranteed point increase of +1 and no transition to either path. Approach decisions took participants down one of the two paths, as chosen by the computer. Crucially, however, there was always a degree of uncertainty as to which of the two paths the participant would transition to if an approach decision was made. The transition probability of each path varied from trial to trial and was explicitly conveyed to the participant at the beginning of each trial. There were five possible sets of probabilities: 10-90%, 30-70%, 50-50%, 70-30%, and 90-10% for transitions to paths 1 and 2, respectively. Once transitioned to a path, the transitions to each room within the sequence were deterministic.

Participants were required to use the value map they had learned in the previous stage, in conjunction with the path transition probabilities presented on each trial, to evaluate the utility of making an approach versus an avoid decision. Optimally, this evaluation would reflect an expected value calculation for both approach and avoid choices, such that:

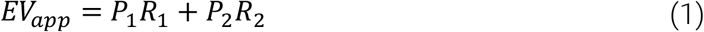

where ***EV***_***app***_ is the expected value of approaching, ***P***_**1**_ and ***P***_**2**_ are the probabilities of transitioning to paths 1 and 2, respectively, and ***R***_**1**_ and ***R*** _**2**_ are the total sums of points for paths 1 and 2, respectively, taking into account the odd rule states (see **Methods, Planning manipulation**). The expected value of avoiding, ***EV***_***AV***_, was always 1. The decision to approach was considered correct if ***EV***_***app***_ **≥ *EV***_***av***_ and the decision to avoid was considered correct if ***EV***_***app***_ **≤ *EV***_***av***_.

After each block, the proportion of correct responses was converted into a monetary value and displayed as a bonus. Participants did not receive feedback on the accuracy of their choices throughout the block. They did, however, observe an animation of their subsequent transitions and change in points (**Fig. 1A**). For “avoid” decisions, a screen was displayed with text stating that they had received 1 point (3 seconds). For “approach” decisions, participants were first shown which path had been selected by the computer according to the transition probability (“Path 1” or “Path 2”, for 3 seconds). Participants were then shown each state within that path one at a time (2 second presentation), underneath which the state value as well as the running total of points collected along the path was displayed. A blank screen was presented between states (randomly jittered duration between 0.5 and 0.8 seconds). A final screen conveyed the total number of points earned for that trial (2 seconds). Trials were separated by a blank screen (1 second).

After an initial practice block, participants completed 6 (behavioural session) or 10 (MEG session) blocks. Each block contained 18 decision trials. In the practice block, participants were given unlimited time to make their decision and did not earn bonus money. In test blocks, participants were given 30 seconds (indicated by an on-screen timer) to make their choice. Responses were disabled for the first 5 seconds to prevent accidental presses and encourage planning. If no response was made after 30 seconds, participants were penalised −1 point and prompted with a warning message (“Too slow!”) and the trial ended.

#### Planning manipulation

A number of additional features were incorporated into the design of the decision trials to encourage planning, as well as to control for certain variables. One feature was what we term the “odd rule”. The purpose of the odd rule was to allow the sum of points along each path to vary from trial to trial, thus encouraging participants to engage in sequential planning. On each trial, the odd rule was applied to two states: one from each path. These two odd rule states were displayed on-screen (as images on forced-choice trials or as text labels on free-choice trials) at the beginning of each trial, alongside the path probabilities (**Fig. 1A**). Participants were instructed that, if the sum of points accumulated up until (and including) an odd rule state was an odd number, then the sign of this cumulative sum would “flip” (i.e., a negative cumulative sum will become positive, and vice versa). This new sum would then be carried over to any subsequent states along the path.

By way of example, assume the values of states A, B, and C in path 1 are −5, −2, and 3, respectively. If state B is the odd rule state, then one must mentally sum the number of points up until (and including) state B (−5 + −2 = −7). One must then consider whether the current sum of points is an odd number. In this case, it is (−7), and thus the sign of the sum is flipped (becoming +7). This value is then carried over to the next state, C (7 + 3 = 10), producing a final outcome of 10. If, instead, state C is the odd rule state, then one sums the number of points up until state C (−5 + −2 + 3 = −4). In this case, the sum of points at the odd-rule state is an even number (−4), and thus no sign-flipping occurs, producing a final outcome of −4. Hence, the final value of each path is entirely dependent on the position of the odd rule state in each path (see **Fig. 1A** for another example). This manipulation increased the variability of final path values across trials.

To further increase the variability in final path values across the experiment, the value of one state from each path changed at the beginning of each block. All state values then remained constant for the duration of the block. So that participants knew which values had changed at the beginning of each block, the first four trials in each block (first six in the practice block) were forced-choice, such that participants could only choose to approach. Forced-choice trials were controlled so that they lead to an equal number of transitions to path 1 and path 2. Any points gained or lost on these trials did not count towards bonus payment and were not included in planning-related MEG analyses, as participants were unable to plan until having observed the updated values in both paths.

Forced-choice trials were also the only trials in which the images were displayed, both during the planning period (where the states with the odd rules were displayed) and the sequence animation. In all other free-choice trials, images were replaced by their text labels (e.g., “cat” or “bicycle”), which had already been shown to participants during the functional localiser (see **Procedure** below). This was done to control for any potential biased visual exposure to the state images during free-choice trials based on choice behaviour (e.g., only deciding to approach when path 1 is more likely) while still periodically reminding participants of the images associated with each room.

Participants were assigned to one of two experimental protocols in a counterbalanced fashion (**Supp. Fig. 1E**). Each protocol was designed to minimise the repetition of odd rule state pairs across trials. These two protocols also captured another feature of the design, in which one path more often resulted in a positive outcome and the other in a negative outcome. This was done to maximise the difference in replay between rewarding and aversive paths, by allowing for some degree of association by repetition. To prevent participants from relying on this consistency (and thus not engaging in sequential planning), 5% of trials were catch trials, where either both paths produced a gain or both produced a loss, thus increasing the utility of planning on every trial. Furthermore, the rewarding and aversive paths swapped positions halfway through the experiment (e.g., if path 1 was consistently rewarding at the beginning, it became consistently aversive, and vice versa for path 2). The starting positions of the rewarding and aversive paths were counterbalanced across the two protocols.

### Procedure

#### Initial session

Participants completed two sessions on consecutive days. The first session was a behavioural-only practice, where participants completed three questionnaires: the 12-item Intolerance of Uncertainty Scale ^61^, 16-item Penn State Worry Questionnaire ^62^, and 30-item Domain-Specific Risk-Taking Scale ^63^, each presented in a random order on a computer (approximately 15 minutes). Participants then completed a shorter 45-minute version of the experiment. The aim of this session was to ensure participants were capable of performing the task (at least 80% performance on the image and value memory tests, and at least 60% correct choices on decision trials) before continuing to the MEG session the following day.

#### Functional localiser

The second session comprised an MEG session. Participants first completed a functional localiser task (30-minutes) and then completed a full 1.5-hour task. In the functional localiser, participants were shown the six unique images (randomly selected per participant) used in the main task. Crucially, these images were different from those shown in the initial behavioural session. On each trial, an image was presented on screen for 1 second (**Supp. Fig. 1B**). After the image disappeared, two words were presented on the left and the right of the screen. One of these was the correct label for the previous image (e.g. “cat”) and the other label was randomly selected from a pool of invalid words. Participants pressed either the left or right button of a 4-button response pad to indicate the correct label. After making a response, the words were replaced by a fixation cross for a randomly jittered inter-trial interval between 0.5 and 1.5 seconds. Correct and incorrect responses produced a green or red cross, respectively. There were four blocks, within which each image was randomly presented 20 times, giving 80 trials in total per image. Across the 26 participants, the mean response accuracy was 97.48% (SD = 2.48%, range = 90.63 to 99.79%).

### MEG analysis

#### MEG acquisition and preprocessing

Participants’ neural activity was measured using a CTF Omega MEG scanner with a 275-channel axial gradiometer whole-head system (CTF Omega, VSM MedTech) at University College London. Participants were seated upright in the scanner and head position was continuously monitored by three head position indicator coils located at the nasion and left and right pre-auricular fiducial points. Data were acquired continuously at 1,200 Hz and participants’ eye movements were recorded using an Eyelink eye-tracking system. Triggers were recorded using a photodiode positioned behind the stimulus presentation screen that detected the onset of a flashing white stimulus (hidden from view) that was synchronised with event onsets.

MEG data from the functional localiser and decision trials were preprocessed using SPM12 (Wellcome Centre for Human Neuroimaging), Fieldtrip (2019), and custom code written in MATLAB R2018b (MathWorks). All code is available on GitHub: https://github.com/jjmcfadyen/approach-avoid-replay. CTF data for each block were imported using OSL (the OHBA Software Library, from OHBA Analysis Group). Trigger onset times and durations were extracted from the photodiode signal and semi-automatically checked for errors. Next, the data were high-pass filtered at 0.5 Hz to reduce slow drift, and a notch filter for 50 Hz was applied to remove line frequency. The data were then downsampled to either 100 Hz (for replay analysis) or 600 Hz (for source reconstruction) to reduce computational load and increase signal to noise ratio. OSL also identified potential bad channels whose characteristics fell outside the normal distribution of values for all sensors.

Independent component analysis was then performed on the data (FastICA, http://research.ics.aalto.fi/ica/fastica), decomposing it into 150 independent spatiotemporal components. Artefactual components were automatically classified using the combined spatial topography, time course, time course kurtosis, and frequency spectrum of all components. For example, eye blink artifacts exhibited high kurtosis (>20), a repeated pattern in the time course, and consistent spatial topographies. The number of excluded components was limited to a maximum of 20. Artefacts were rejected by subtracting them out of the data. All subsequent analyses were performed directly on the filtered, cleaned MEG signal, in units of femtotesla.

The data were then divided into different epochs using the trigger onsets and durations. For the functional localiser, epochs were created for the image onset (−0.1 to 0.8 seconds post-stimulus onset). For the decision trials in the main task, epochs were created for the planning time (−0.1 seconds before trial onset to the response time). Artefactual sensors identified by OSL were interpolated for all epochs, and artefactual functional localiser trials were excluded from the classification procedure.

#### Image classification

We used Temporal Delayed Linear Modelling (TDLM) to characterise patterns of neural dynamics during the task ^64^, as performed in previous studies ^12,14,15,17,44^. First, for each participant, we classified patterns of multivariate neural activity evoked by each image in the functional localiser (**Fig. 2A**). We selected data from 0 to 300 ms from each functional localiser epoch, excluding incorrect and artefactual trials, as well as trials where response time was > 5 standard deviations from the mean per participant (average of 78 trials per stimulus, per participant; SD = 2, range = 73 to 80). We then constructed a series of Lasso-regularised logistic regression models. Each model received data from a single time sample (0 to 300 ms, at 10-ms resolution) across all trials. Hence, we constructed separate models (per time sample, and per image; 31 × 6) per participant, each using a trials × sensors (e.g., 480 × 275) data matrix and a binary vector indicating which trials belonged to that image. For each model, we appended a duplicate-sized matrix of zeros to the data matrix to reduce the spatial correlation between each model.

Each lasso-regularised logistic regression model used a range of 100 regularisation parameters (λ) sampled from a half-Cauchy distribution (γ = 0.05, range = 0.0001 to 1). Thus, each model produced a λ × sensors (100 × up to 275) matrix of slope coefficients (**Fig. 2B**), as well as a vector of intercept coefficients for each λ. We refer to these coefficients as our binomial classifiers, each of which are trained to distinguish the sensor data associated with one image as compared to all other images.

To evaluate the accuracy of each classifier per participant, we conducted a *K*-folds cross-validation procedure. *K* was set to the minimum number of trials per stimulus for that participant. In each fold, a test set was created by randomly taking one sample from one exemplar trial per stimulus. The remaining data was used for training. Random selection of the test data was controlled to maximise equal sampling across trials. The classifiers per state generated from the training dataset were then applied to the six test trials (one for each stimulus). Thus, for a given fold, a score of 1 or 0 was given for whether each state classifier maximally predicted the correct trial. The accuracy of each state classifier was given by the average score across folds.

For each subject, we selected λ that produced the highest mean accuracy across state classifiers (λ: M = 0.0017, SD = 0.0015). We then averaged the classification accuracy across states per subject and examined which training times produced the highest accuracy across subjects (**Fig. 2C**). Overall average state classification accuracy exceeded chance (16.66%) for all subjects from 80 ms onwards, peaking at 120 ms (48.97%). Classifier training times from 110 to 150 ms made up the top 15% performance (all > 45.80% accuracy).

#### Sequential state reactivation

Using our state classifiers, we then estimated the degree to which images were sequentially reactivated in the brain while participants planned whether to approach or avoid the state space in each trial. We utilised an updated general linear modelling approach, which encapsulates a lagged cross-correlation between the evidence for state-to-state transitions. This method produces an overall “sequenceness” statistic at different time intervals, or “lags”. We employed this approach on a trial-by-trial basis per participant, using neural data collected during the planning period.

In the first step, we estimated the degree to which each state was reactivated during the planning period of free-choice decision trials by multiplying the spatiotemporal MEG data by each state classifier’s beta estimates. We used state classifiers trained at 120 ms post-stimulus onset, which had the highest cross-validated accuracy across subjects. We then entered the resultant time series of predicted state reactivation (states × time matrix; **Fig. 2D**) per trial into a 2-level general linear model designed to test whether reactivation of each stimulus occurred in a specific order at different time intervals.

At the first level, we performed a family of multiple regressions for each state’s reactivation time series (***i ϵ* [1: 6]**), in which a time-lagged copy of the reactivation time series for state ***j*** (***X*(*Δt*)**_***j***_) predicts the original, unshifted reactivation time series of state ***i*** (***X***_***i***_). The time lags ranged from 0 to 600 ms, in 10 ms bins. Hence, this analysis evaluated the average likelihood that stimulus ***i*** is followed by stimulus ***j*** after a time lag of ***Δt***. Separate linear models were estimated for each stimulus ***i*** and each time lag ***Δt*:**

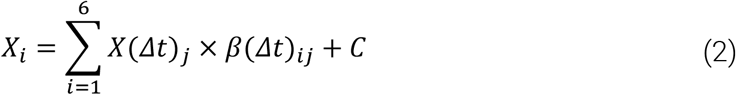

where *C* is a constant term and ***β*(*Δt*)**_***ij***_ is a coefficient derived from ordinary least-squares that captures the unique influence of ***X***_***i***_ on ***X*(*Δt*)**_***j***_. These coefficients are then used to form 6 × 6 empirical transition matrices, ***B*(*Δt*)**, for each time lag.

At the second level, we quantified the evidence for specific, hypothesised state-to-state transitions. In this task, the key state-to-state transitions were A → B and B → C (path 1), as well as D → E and E → F (path 2). These transitions were declared by separate 6 × 6 binary matrices for hypothesised forward (***T***_***F***_) and backward (*T*_*B*_) transitions, where ***T***_***F***_ = ***T***_***B***_. The evidence for the hypothesised transitions was then quantified by:

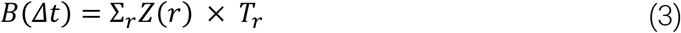

where ***r*** is the total number of all regressors included in the second level. These regressors included ***T***_***F***_, ***T***_***B***_, ***T***_***auto***_ (an identity matrix of self-transitions to control for autocorrelation), and ***T***_***const***_ (a constant matrix that models away the average of all transitions, ensuring that any weight on ***T***_***F***_ and ***T***_***B***_ was not due to general dynamics in background neural dynamics). Note that there were four versions of ***T***_***F***_ and ***T***_***B***_, one for each hypothesised transition (A → B, B → C, D → E, and E → F). This allowed us to examine the evidence of replay of each transition specifically, which was critical to our path-specific analyses. ***Z*** is the weight for each regressor, representing the evidence for the hypothesised state-to-state transitions. ***Z***_***F***_ and ***Z***_***B***_ are evidence for forward and backward transitions, respectively. A forwards-minus-backwards sequenceness measure, ***Z***_***D***_, was also computed by performing ***Z***_***F***_ **− *Z***_***B***_, thus removing common variance. Repeating equation 3 at each time lag produces a time series of sequenceness at different intervals, where smaller intervals indicate more time-compressed replay (**Fig. 2E**)

To determine the statistical significance of ***Z*** (averaged across the four transitions and all trials per participant), we employed non-parametric permutation testing at the second level. We generated a null distribution by generating all possible invalid versions of ***T***_***F***_ and ***T***_***B***_, such that they only included cross-path transitions (e.g., A to E, B to D, etc.). This produced 40 null versions of ***Z***. We then calculated a significance threshold for our valid ***Z*** by taking the maximum absolute value of each null ***Z*** and computing the 95th percentile for ***Z***_***F***_ and ***Z***_***B***_ (one-sided test) or the 2.5th and 97.5th percentile for ***Z***_***D***_(two-sided test). Thus, values of ***Z*** were deemed statistically significant (FWE < 0.05) if they exceeded these significance thresholds.

To account for inter-subject variability in classification accuracy across training times and their relevance to replay, we also computed sequenceness using classifiers trained on 110 ms and 130 ms (10 ms either side of the winning training time). Thus, we computed sequenceness three times per subject, and chose the classifier training time (110 ms, 120 ms, or 130 ms) that produced the greatest absolute value of ***Z***_***D***_ across lags, averaged across all transitions (110 ms = 11 subjects, 120 ms = 10 subjects, 130 ms = 5 subjects).

#### Source localisation

To investigate the neural sources underlying replay during planning, we used a procedure for identifying replay onsets performed in previous studies ^14,15,44^. We first normalised state reactivation probabilities per time sample, and then multiplied the normalised reactivation time series for state ***i* (*X***_***i***_) by the that of the following state ***j*** at time lag ***t* (*X*(*Δt*)**_***j***_). We computed this separately for time lags 20 to 90 ms, as each of these were found to be significant for forwards replay on average across all transitions (Fig. 3E), and chose the time lag that produced the highest combined reactivation at each time point. This produced time series of reactivation likelihood, ***R***_***ij***_, for each transition, per trial.

The above time series were used to determine the onset of replay events throughout the planning period of each decision trial. To do this, ***R***_***ij***_ was thresholded at its 95^th^ percentile to only include high-magnitude putative replay onset events within each trial. We also imposed a constraint that a replay onset event must be preceded by 100 ms of replay-onset-free time.

We epoched the MEG data according to the replay onsets (−100 to 150 ms surrounding replay onset) and baseline corrected the data using a −100 and −50 ms window. We then transformed these data to a three-dimensional grid in MNI space (grid step = 5 mm) using a linearly constrained minimum variance beamformer ^65,66^, as implemented in OSL. Forward models were generated on the basis of a single shell using superposition of basis functions that approximately corresponded to the plane tangential to the MEG sensor array. The sensor covariance matrix for beamforming was estimated using data in a broadband (1 to 150 Hz) frequency range.

At the first level, we computed one-sample tests on whole-brain source activity at each time point using nonparametric permutation testing ^67^ as implemented in OSL. First-level contrasts were defined as either any replay event, or as the difference between rewarding and punishing path replay (see **Supplementary materials**). We selected the resultant t-maps for each participant and smoothed the images in SPM12 using an 8 mm FWHM Gaussian kernel. We then entered these into one-sample t-tests (for either any replay event or for the difference between rewarding and punishing path replay events) in SPM12 for group-level inference. All statistics are p < 0.05, FWE-corrected at the cluster level. We also investigated the hippocampus specifically by using an initial threshold of p < 0.001 uncorrected and then applying a small-volume correction using an anatomical mask (created by automatic subcortical segmentation Freesurfer using an MNI template; http://surfer.nmr.mgh.harvard.edu/).

### Multi-level modelling

All analyses were conducted on the MEG session, as the initial behavioural session served purely to acquaint participants with the task structure. We adopted a multi-level modelling approach, which allowed us to examine effects on a trial-by-trial basis. This approach also allowed us to compare conditions with unbalanced trial numbers (e.g., “approach” decisions mostly consisted of trials where reward probability was high, and vice versa for “avoid” decisions).

We used the lme4 package implemented in R v3.6. We constructed a series of models that either used: a) choice as a binomial dependent variable, or b) sequenceness as a linear dependent variable. In all models, forced choice trials and catch trials (i.e., trials where both paths resulted in an overall loss or both resulted in an overall gain) were excluded. All predictors were mean-centred. To ensure convergence, the bobyqa optimiser was used and set to 10^6^ iterations. Significant interaction terms were followed up by simple slopes analyses using the “interactions” package in R, FDR-corrected for multiple comparisons, and the “emmeans” package in R. We also ensured that all models produced a variable inflation factor (VIF) below 5 and that autocorrelation within the residuals of each model was minimal, as assessed by a Durbin-Watson test ^68^; see **Supplementary materials**).

For models including individual differences, we used principal components analysis to reduce the dimensionality of the three self-report questionnaires (intolerance of uncertainty, worry, and risk-taking across 7 domains: ethical, social, health, financial, and recreational) completed at the beginning of the behavioural session. We identified two principal components that together explained 60.55% of the variance (41.52% and 29.49%, respectively; eigenvalues = 1.548, 1.357, 1.040, 0.944, 0.684, 0.457, 0.350). The first component mapped positively on to risk-taking questionnaire scores, while the second component mapped negatively on to intolerance of uncertainty and worry. We refer to these two components as risk-seeking and anxiety, respectively. For interpretability, we inverted these factors, such that more positive values represented higher risk-aversion and higher anxiety, respectively.

