## Supplementary materials for "Differential replay for reward and punishment paths predicts approach and avoidance"

#### Figures

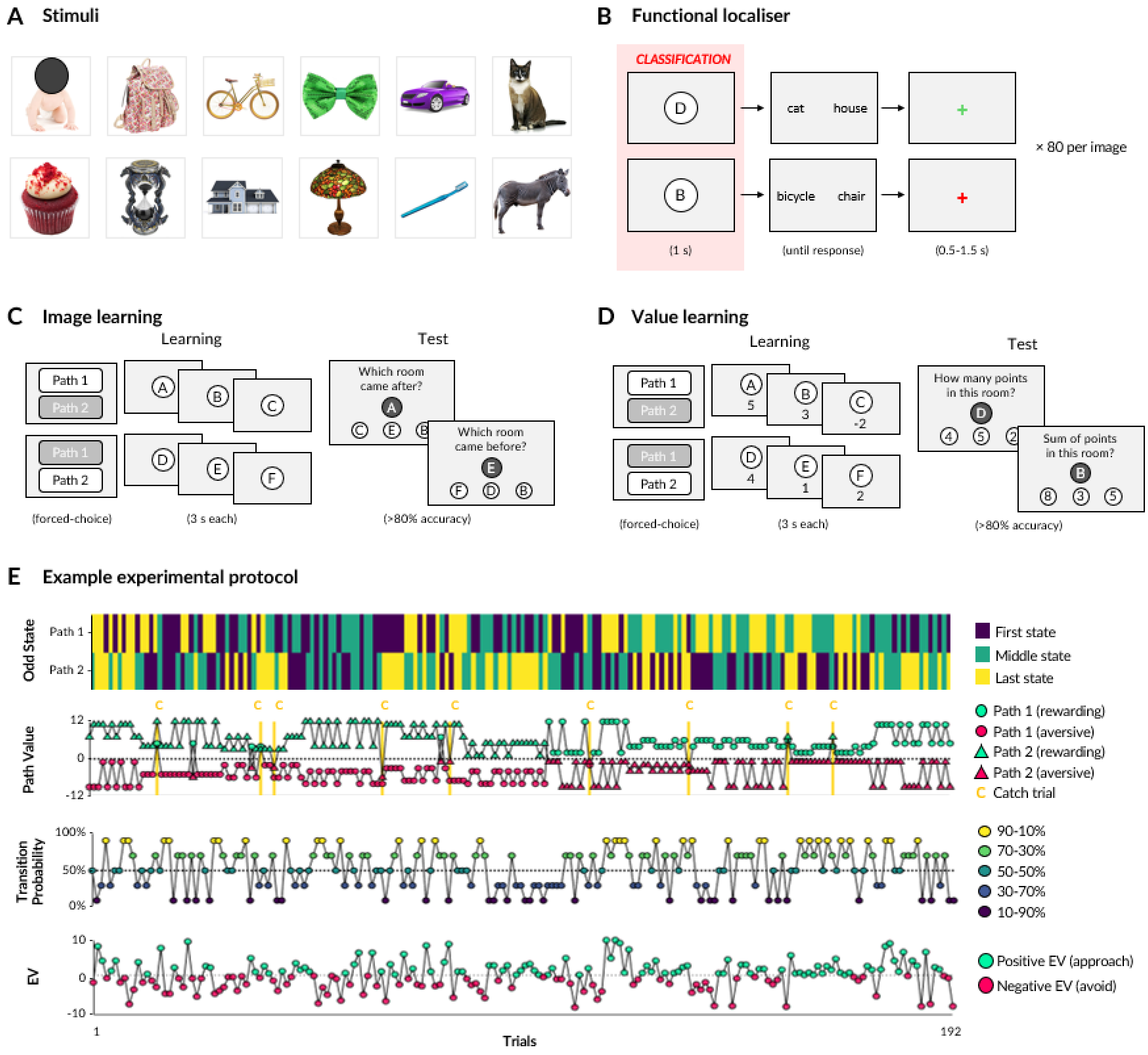


**Supplementary Figure 1. Experimental paradigm. (A)** All 12 stimuli used in the experiment. Six stimuli were pseudo-randomly allocated to each participant, per session, ensuring equal allocation of each stimulus. Note that a face has been obscured for compliance with *bioRxiv* policy. **(B)** For visualisation purposes, stimuli are represented by letters: A, B, and C for path 1, and D, E, and F for path 2. In an initial functional localiser in session 2, participants viewed an image and then reported the correct label (left or right). Correct and incorrect responses produced a green or red fixation cross, respectively. **(C)** To learn the image order, participants selected either Path 1 or Path 2 (forced-choice, 2 selections of each) and then observed an animation of the sequence of states along each path. Participants were then tested on their memory for the image order. (D) After image learning, participants observed the animated sequences again but this time with the value of each state displayed underneath each image. Participants were tested on their memory for the values associated with each state, as well as their ability to calculate the cumulative sum at different states. (E) Two protocols were used, counterbalanced across participants. One such protocol is shown. The designation of the odd rule to one state per path (first row) dictated the final value of each path (second row). One path was mostly negative (here, starting with path 1; circles) and the other mostly positive (here, starting with path 2; triangles), with the exception of catch trials (marked in yellow). This tendency switched halfway through the experiment. The transition probability (third row) to each path, in combination with the path value, dictated the expected value of approaching (fourth row). Expected value was a sum of each path's value weighted by its probability. When the expected value was greater than 1 (which was guaranteed outcome of avoiding), participants should approach (green); otherwise, participants should avoid (red).

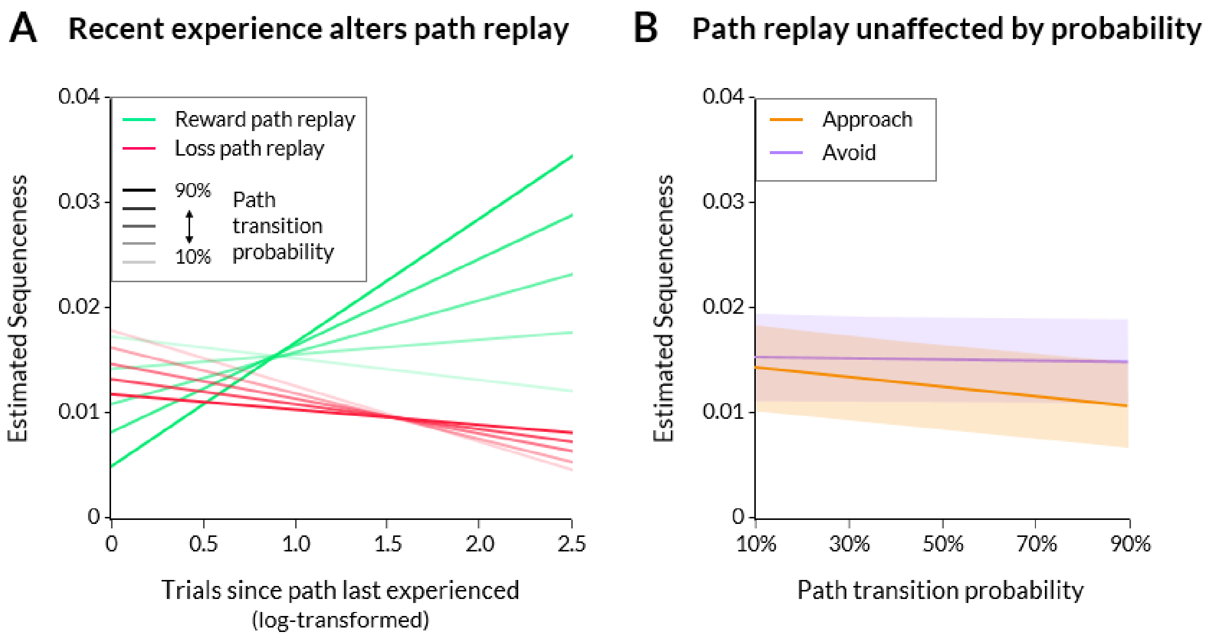


Supplementary Figure 2. Modulation of path replay by experience and expectations. (A) Replay strength (y axis) during planning as predicted by a model containing path type (reward or loss), path experience (x axis), and path transition probability (darker lines indicate higher transition probability). Evidence of rewarding path replay increased when rewarding paths had not been experienced for longer, whereas the opposite was true for punishing paths. This was most prominent when rewarding paths were more likely to be transitioned to. (B) Replay strength (y axis) during planning was not significantly predicted by a model containing path probability (x axis) and choice (approach or avoid).

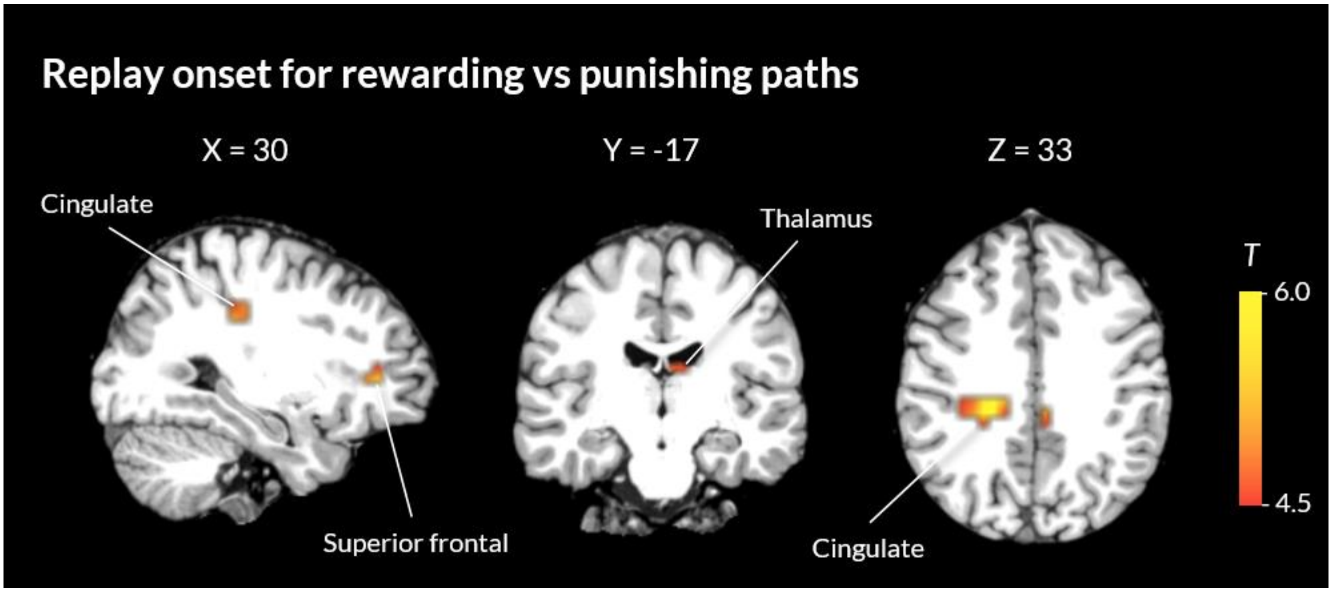


Supplementary Figure 3. Beamforming analysis on replay onsets. A contrast between replay events for rewarding vs punishing paths showed significant sources in the cingulate cortex, superior frontal gyrus, and thalamus. Clusters are thresholded at $p < 0.05$, whole brain FWE-corrected.

We hypothesised that replay events during planning might be generated by different sources depending on whether the replayed transition belonged to a reward or loss path. We found significant clusters in the cingulate cortex, as well as the superior frontal gyrus and thalamus (whole brain, $p < 0.05$ FWE-corrected), which were all more strongly activated at the onset of reward path replay events than loss path replay events.

We then compared source reconstructions of replay events (irrespective of reward or loss) preceding decision to approach vs avoid. We found significantly greater activation in the cingulate cortex, thalamus, and precentral gyrus for replay events that occurred while participants planned to avoid compared to approach (whole brain, $p < 0.05$ FWE-corrected). This closely aligns with the neural regions that were more activated for reward than loss path replay. We also contrasted the difference between reward and loss path replay between choice types, but did not observe any significant effects.

### Model output tables

Model 1: Choice ~ (Reward magnitude × Loss magnitude × Transition probability) + Certainty + RT + (1 | Subject)

| Fixed Effect | ß | SEM | p | |
|---|---|---|---|---|
| (Intercept) | 0.03 | 0.17 | 0.852 | |
| Reward magnitude | 0.11 | 0.02 | < 0.001 | *** |
| Loss magnitude | 0.05 | 0.02 | 0.011 | * |
| Transition probability | 6.46 | 0.25 | < 0.001 | *** |
| Certainty | 0.32 | 0.07 | < 0.001 | *** |
| RT | 0.01 | 0.01 | 0.477 | |
| Reward magnitude × Loss magnitude | 0.01 | 0.01 | 0.202 | |
| Reward magnitude × Transition probability | 0.50 | 0.09 | < 0.001 | *** |
| Loss magnitude × Transition probability | -0.03 | 0.09 | 0.744 | |
| Reward magnitude × Loss magnitude × Transition probability | 0.02 | 0.03 | 0.464 | |

*Variable inflation factor = 1.03 to 1.55, Durbin-Watson = 1.86*

Model 2: Sequenceness ~ (Replay type × Choice) + RT + (1 | Subject/Lag)

| *Fixed Effect* | *ß* | *SEM* | *p* | |
|---|---|---|---|---|
| (Intercept) | 0.01 | 0.00 | 0.009 | ** |
| Replay type | 0.01 | 0.00 | < 0.001 | *** |
| Choice | 0.00 | 0.00 | 0.309 | |
| RT | -0.00 | 0.00 | 0.008 | ** |
| Replay type × Choice | -0.01 | 0.00 | < 0.001 | *** |

*Variable inflation factor = 1.01 to 3.46, Durbin-Watson = 1.76*

Model 3: Sequenceness ~ (Recency × Replay type × Transition probability) + RT + (1 | Subject/Lag)

| *Fixed Effect* | *ß* | *SEM* | *p* | |
|---|---|---|---|---|
| (Intercept) | 0.01 | 0.00 | 0.006 | ** |
| Recency | -0.00 | 0.00 | 0.001 | *** |
| Replay type | 0.00 | 0.00 | 0.009 | ** |
| Transition probability | -0.00 | 0.00 | 0.049 | * |
| RT | -0.00 | 0.00 | 0.01 | ** |
| Recency × Replay type | 0.01 | 0.00 | < 0.001 | *** |
| Recency × Transition probability | 0.01 | 0.00 | 0.155 | |
| Replay type × Transition probability | 0.00 | 0.00 | 0.74 | |
| Recency × Replay type × Transition probability | 0.01 | 0.01 | 0.01 | ** |

*Variable inflation factor = 1.02 to 2.05, Durbin-Watson = 1.76*

Model 4: Sequenceness ~ (Transition probability × Choice) + RT + (1 | Subject/Lag)

| *Fixed Effect* | *ß* | *SEM* | *p* | |
|---|---|---|---|---|
| (Intercept) | 0.01 | 0.00 | 0.001 | *** |
| Transition probability | -0.00 | 0.00 | 0.83 | |
| Choice | -0.00 | 0.00 | 0.003 | ** |
| RT | -0.00 | 0.00 | 0.008 | ** |
| Transition probability × Choice | -0.00 | 0.00 | 0.19 | |

*Variable inflation factor = 1.01 to 2.49, Durbin-Watson = 1.76*

Model 5: Choice ~ (Expected value × Differential replay) + Certainty + RT + (1 | Subject/Lag)

| *Fixed Effect* | *ß* | *SEM* | *p* | |
|---|---|---|---|---|
| (Intercept) | -0.04 | 0.10 | 0.735 | |
| Expected value | 0.43 | 0.01 | < 0.001 | *** |
| Differential replay | -0.71 | 0.13 | < 0.001 | *** |
| Certainty | 0.27 | 0.02 | < 0.001 | *** |
| RT | -0.00 | 0.00 | 0.646 | |
| Expected value × Differential replay | 0.13 | 0.05 | 0.008 | ** |

*Variable inflation factor = 1.02 to 1.05, Durbin-Watson = 1.87*

Model 6: Choice ~ (Expected value × Rewarding path replay) + (Expected value × Punishing path replay) + Certainty + RT + (1 | Subject/Lag)

| *Fixed Effect* | *ß* | *SEM* | *p* | |
|---|---|---|---|---|
| (Intercept) | -0.04 | 0.10 | 0.735 | |
| Expected value | 0.43 | 0.01 | < 0.001 | *** |
| Rewarding path replay | -1.23 | 0.19 | < 0.001 | *** |
| Punishing path replay | 0.19 | 0.19 | 0.313 | |
| Certainty | 0.27 | 0.02 | < 0.001 | *** |
| RT | -0.00 | 0.00 | 0.598 | |
| Expected value × Rewarding path replay | 0.12 | 0.07 | 0.09 | |
| Expected value × Punishing path replay | -0.15 | 0.07 | 0.031 | * |

*Variable inflation factor = 1.01 to 1.05, Durbin-Watson = 1.87*

Model 7: Choice ~ (Expected value × Differential replay × Risk-aversion) + (Expected value × Differential replay × Anxiety) + Certainty + (1 | Subject/Lag)

| *Fixed Effect* | *ß* | *SEM* | *p* | |
|---|---|---|---|---|
| (Intercept) | -0.03 | 0.10 | 0.745 | |
| Expected value | 0.43 | 0.01 | < 0.001 | *** |
| Differential replay | -0.62 | 0.14 | < 0.001 | *** |
| Risk-aversion | 0.03 | 0.06 | 0.644 | |
| Anxiety | -0.07 | 0.07 | 0.284 | |
| Certainty | 0.27 | 0.02 | < 0.001 | *** |
| Expected value × Differential replay | 0.11 | 0.05 | 0.028 | * |
| Expected value × Risk-aversion | -0.01 | 0.00 | 0.048 | * |
| Differential replay × Risk-aversion | 0.38 | 0.09 | < 0.001 | *** |
| Expected value × Anxiety | 0.01 | 0.00 | 0.168 | |
| Differential replay × Anxiety | 0.31 | 0.11 | 0.003 | ** |
| Expected value × Differential replay × Risk-aversion | 0.02 | 0.03 | 0.54 | |
| Expected value × Differential replay × Anxiety | -0.10 | 0.04 | 0.014 | * |

*Variable inflation factor = 1 to 1.12, Durbin-Watson = 1.87*

Model 8: Choice ~ (Expected value × Rewarding path replay × Risk-aversion) + (Expected value × Rewarding path replay × Anxiety) + (Expected value × Punishing path replay × Risk-aversion) + (Expected value × Punishing path replay × Anxiety) + Certainty + (1 | Subject/Lag)

| *Fixed Effect* | *ß* | *SEM* | *p* | |
|---|---|---|---|---|
| (Intercept) | -0.04 | 0.10 | 0.725 | |
| Expected value | 0.44 | 0.01 | < 0.001 | *** |
| Rewarding path replay | -1.14 | 0.19 | < 0.001 | *** |
| Risk-aversion | 0.02 | 0.06 | 0.705 | |
| Anxiety | -0.08 | 0.07 | 0.245 | |
| Punishing path replay | 0.05 | 0.19 | 0.794 | |
| Certainty | 0.27 | 0.02 | < 0.001 | *** |
| Expected value × Rewarding path replay | 0.12 | 0.07 | 0.102 | |
| Expected value × Risk-aversion | -0.01 | 0.00 | 0.116 | |
| Rewarding path replay × Risk-aversion | 0.22 | 0.13 | 0.084 | |
| Expected value × Anxiety | 0.01 | 0.00 | 0.134 | |
| Rewarding path replay × Anxiety | 0.20 | 0.15 | 0.189 | |
| Expected value × Punishing path replay | -0.11 | 0.07 | 0.13 | |
| Risk-aversion × Punishing path replay | -0.47 | 0.12 | < 0.001 | *** |
| Anxiety × Punishing path replay | -0.45 | 0.15 | 0.003 | ** |
| Expected value × Rewarding path replay × Risk-aversion | 0.13 | 0.05 | 0.004 | ** |
| Expected value × Rewarding path replay × Anxiety | -0.01 | 0.06 | 0.85 | |
| Expected value × Risk-aversion × Punishing path replay | 0.08 | 0.04 | 0.059 | |
| Expected value × Anxiety × Punishing path replay | 0.20 | 0.05 | < 0.001 | *** |

*Variable inflation factor = 1 to 1.16, Durbin-Watson = 1.87*

Comparison of models containing risk-aversion and/or anxiety as predictors of choice

| *Model definitions* | |
|---|---|
| Model A | Choice ~ (EV × Replay_differential) + Certainty + RT + (1 \| Subject/Lag) |
| Model B | Choice ~ (EV × Replay_differential) + Risk aversion + Certainty + (1 \| Subject/Lag) |
| Model C | Choice ~ (EV × Replay_differential) + Anxiety + Certainty + (1 \| Subject/Lag) |
| Model D | Choice ~ (EV × Replay_differential) + Risk aversion + Anxiety + Certainty + (1 \| Subject/Lag) |
| Model E | Choice ~ (EV × Replay_differential × Risk aversion) + Certainty + (1 \| Subject/Lag) |
| Model F | Choice ~ (EV × Replay_differential × Anxiety) + Certainty + (1 \| Subject/Lag) |
| Model G | Choice ~ (EV × Replay_differential × Risk aversion) + Anxiety + Certainty + (1 \| Subject/Lag) |
| Model H | Choice ~ (EV × Replay_differential × Anxiety) + Risk aversion + Certainty +(1 \| Subject/Lag) |
| Model I | Choice ~ (EV × Replay_differential × Risk aversion) + (EV × Replay_differential × Anxiety) + Certainty + (1 \| Subject/Lag) |

| *Comparison* | $\chi^2$ | *p* |
|---|---|---|
| B vs A | 0.122 | < 0.001 *** |
| C vs B | 0.900 | < 0.001 *** |
| D vs C | 0.352 | 0.553 |
| E vs D | 19.249 | < 0.001 *** |
| G vs E | 0 | 1 |
| F vs G | 3.172 | 0.075 |
| H vs F | 0 | 1 |
| I vs H | 22.031 | < 0.001 *** |
